# Quantifying Visual Acuity for Pre-Clinical Testing of Visual Prostheses

**DOI:** 10.1101/2022.05.16.492214

**Authors:** Martin Spencer, Tatiana Kameneva, David B. Grayden, Anthony N. Burkitt, Hamish Meffin

## Abstract

Visual prostheses currently restore only limited vision. More research and pre-clinical work are required to improve the devices and stimulation strategies that are used to induce neural activity that results in visual perception. Evaluation of candidate strategies and devices requires an objective way to convert measured and modelled patterns of neural activity into a quantitative measure of visual acuity.

This study presents an approach that compares evoked patterns of neural activation with target and reference patterns. A d-prime measure of discriminability determines whether the evoked neural activation pattern is sufficient to discriminate between the target and reference patterns and thus provide a quantified level of visual perception in the clinical Snellen and MAR scales. The measure was accurate in providing an estimate of the perceivable feature sizes in scaled standardized “C” and “E” optotypes.

The approach was used to assess the visual acuity provided by two alternative stimulation strategies applied to simulated retinal implants with different phosphene sizes and electrode pitch configurations. It was found that when there is substantial overlap in neural activity generated by different electrodes, an estimate of acuity based only upon electrode pitch is incorrect; our proposed method gives an accurate result in these circumstances.

Quantification of visual acuity using this approach in pre-clinical development will allow for more rapid and accurate prototyping of improved devices and neural stimulation strategies.

## 1 Introduction

Visual prostheses have been developed to provide vision for those with untreatable degenerative visual conditions [1–3]. Electrical stimulation via an electrode activates neurons along the visual pathway that, in turn, creates visual perception in the form of phosphenes [4]. Retinal implants are proposed as a treatments for diseases and conditions, such as Retinitis Pigmentosa and Macular Degeneration, that damage the photoreceptors of the retina but largely leave the remaining retina intact [5,6]. Cortical implants are proposed as treatments for conditions that cause damage to the retina and optic nerve [7].

While several different types of retinal implants have been implanted previously, either through clinical trials, or commercially, no clinically approved devices are currently being manufactured, partly because the level of visual acuity they provide is limited [8]. Development of implants via improvements in vision processing strategies, stimulation strategies and hardware will require pre-clinical development using computational models and *in vitro* and *in vivo* experimentation.

A common assumption is that the visual acuity provided by an implant is determined by the electrode pitch (the spacing between electrodes, figure 4 in [1], figure 1 in [4], [9]). Under this assumption, increasing visual acuity can be achieved by increasing the number and density of electrodes. However, this assumption is flawed since the spread of current and thus the spread of neural activation from each electrode does not usually change with denser electrodes. Therefore, when the pitch is decreased, neural activity, and therefore perceived phosphenes, increasingly overlap. This results in blurred perception and low visual acuity with simultaneous stimulation or, with sequential stimulation, redundant electrodes that do not contribute to forming images [10].

**Figure 1:**
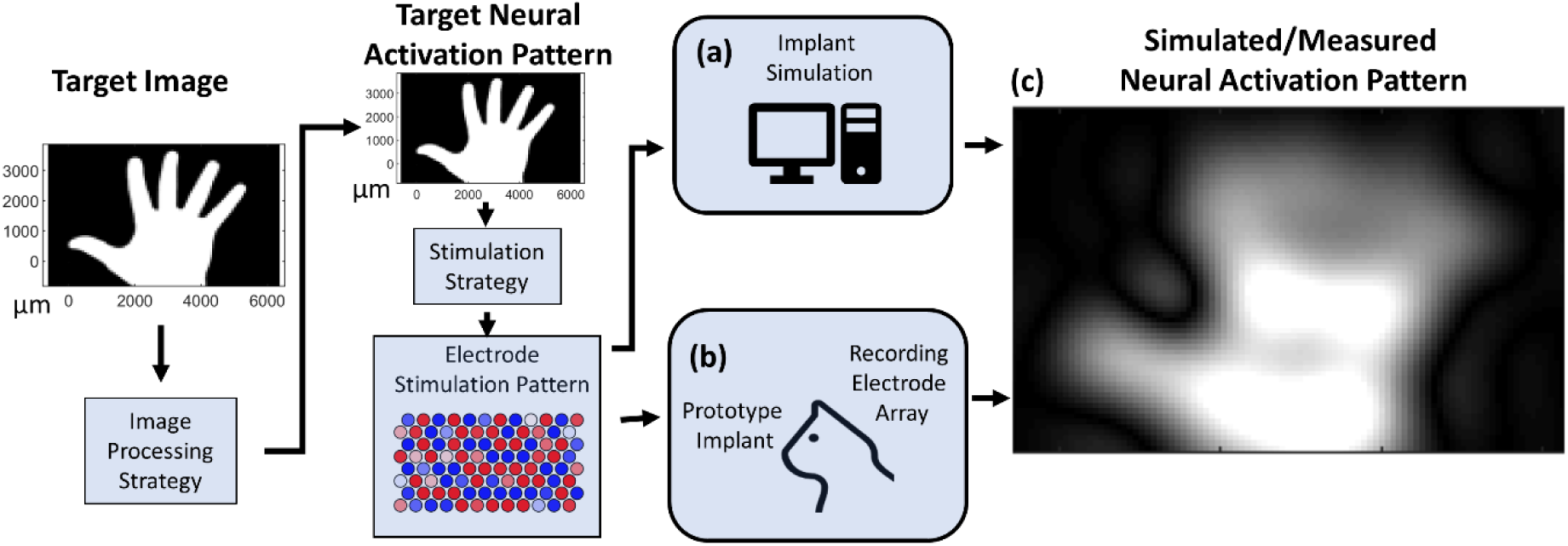
Pre-clinical measures of neural activation in response to electrical stimulation. A target image is processed to create a target neural activation pattern. Then a stimulation strategy determines the electrode simulation pattern that aims to induce the target neural activation pattern in the neural tissue. (a) A computational model of neural activation. (b) An experimental animal model of neural activation using stimulating and recording arrays. (c) Resulting simulated or measured neural activation patterns from either (a) or (b).

Computational and animal models of prosthetic vision can simulate or measure patterns of neural activation in response to stimulation by an implant (Figure 1). Throughout this investigation we assume that neural activity is closely related to visual perception and that spreads of neural activity are closely associated with the spreads of visual perception referred to as phosphenes. In computational models, this pattern is simulated with a map of neural activation made up of pixels representing local neural activation levels (Figure 1a) [11,12]. In animal models, neural activation can be measured via recording electrodes located proximal to the tissue of the retina or cortex [13,14]. The electrical signal from each channel is processed to remove noise and artifacts and the spike (action potential) counts or local field potentials are obtained as measures of neural activation (Figure 1b). In each case, the models produce spatio-temporal patterns of neural activation (Figure 1c) that can be compared to intended target patterns of neural activation using a metric such as the mean squared error (MSE) difference. This metric computes the mean of the square of the difference in amplitudes of all the pixels. This allows different outcomes to be compared and defines a useful objective function for mathematical optimization [15]. However, this does not provide an accurate estimation of *clinical visual acuity* provided by a particular implant configuration and stimulation strategy.

We propose a method to quantify *clinical visual acuity* from these pre-clinical measurements of neural activity patterns.

In a clinical context, it is possible to perform tests of visual acuity based on a person’s ability to distinguish between different letters, called optotypes. These measurements take the form of responses to questions about the identities or orientations of these optotypes. This results in a measure of visual acuity as the Snellen ratio, decimal acuity, minimum angle of resolution in arcminutes (MAR), or logMAR (the log of the MAR value). Decimal acuity is the decimal value of the Snellen ratio and MAR is the inverse of decimal acuity. In cases of very low vision, visual perception is tested in a less quantitative fashion using measures such as a participant’s ability to count fingers, perceive hand motion, and/or perceive light [16]. The participant might also be asked to distinguish between the orientations of sinusoidal gratings or the locations of spots of different sizes [16].

Clinical measurements all require the participant to cognitively assess their perception and answer questions about what they see. In a pre-clinical context, however, this cognitive assessment is not possible. A standard method to convert patterns of neural activation into a clinical measure of visual acuity does not currently exist.

In this study, we propose a method to convert patterns of neural activity measured *in vitro, in vivo* or in simulations into a Snellen measure of visual acuity (which is easily converted into a decimal acuity value or a MAR or logMAR value). This process of acuity assessment is proposed to objectively assess implants in pre-clinical settings, thereby accelerating the development of devices and stimulation strategies that will have improved clinical outcomes.

## 2 Methods

### 2.1 Acuity Measurement

Our proposed acuity measurement is devised to replicate the clinical process of asking a participant to accurately distinguish between spots at different positions and gratings of different orientations. A spot of a particular size and position or a grating of a particular period, orientation, and phase are separately created as different targets of neural activity pattern (Figure 2a). A spot is created as a circularly symmetric half-period sinusoid, so that the spot and grating feature sizes are equivalent and can be quantified in cycles/mm across the neural tissue of the retina. It is important to use these two alternatives both to replicate the proess undertaken clinically and also because the two approaches give somewhat different results when acuity is limited by the size of the array or the pitch of the electrodes.

**Figure 2:**
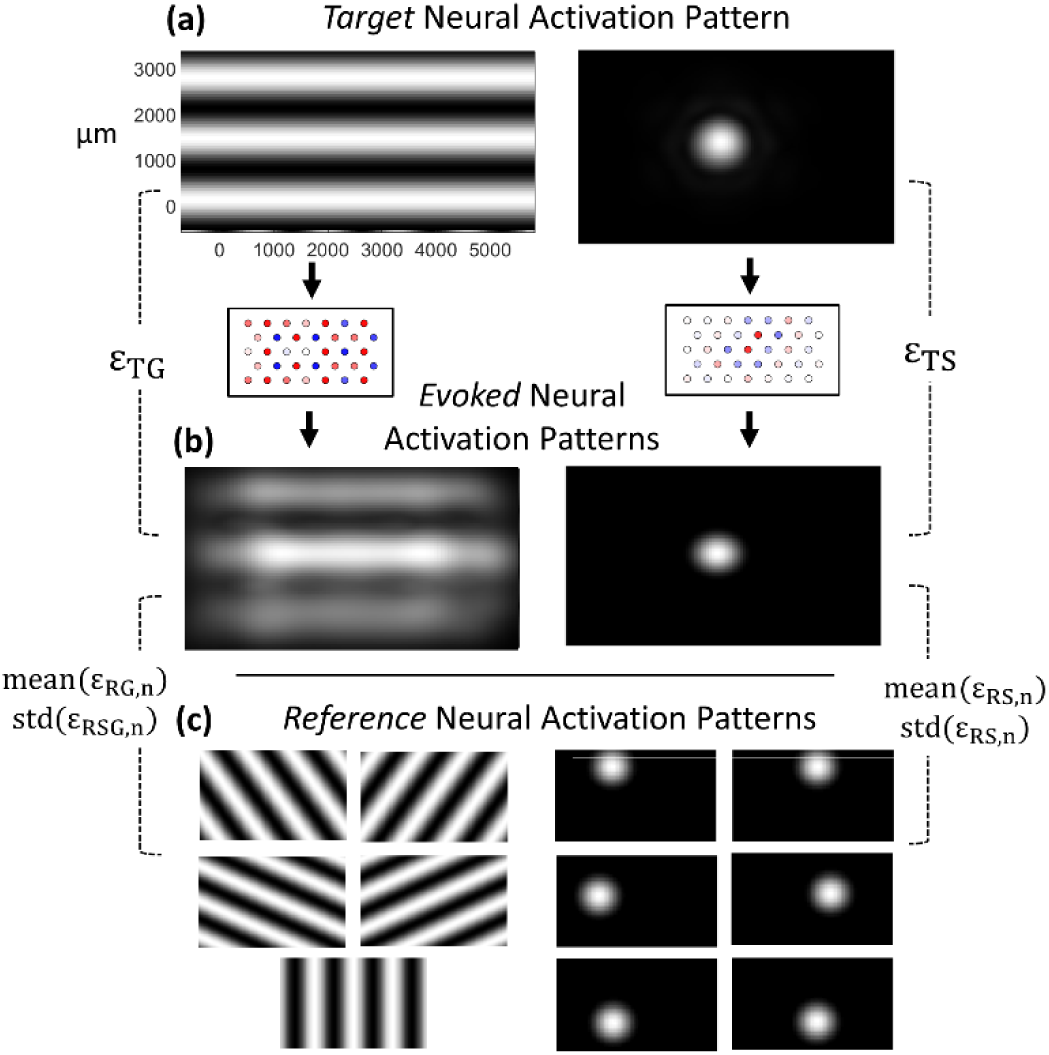
Schematic of the acuity measurement procedure utilising measured or simulated patterns of neural activation. (a) Spot and grating target patterns of a particular feature size are converted into patterns of electrode stimulation using a stimulation strategy. (b) Via either computation or experiment, the electrical stimulations are converted into evoked neural activity patterns. The evoked patterns are used to calculate mean squared error differences for spot (ε_TS_) and grating (ε_TG_) target patterns as well as (c) a set of five grating (ε_RS,n_) and six spot (ε_RG,n_) reference patterns that are used to compute means and standard deviations.

In this investigation, we assume that a visual field of approximately 1° corresponds to a retinal distance of 288 μm. This is based on published experimental observation [17]. Using this value, feature size in cycles/mm can be converted to an acuity measure using the fact that, for human vision, 20/20 vision is equivalent to a spatial frequency of roughly 100 cycles per mm on the retina [18]. This scales linearly such that 20/200 and 20/2000 vision are equivalent to approximately 10 cycles/mm and 1 cycle/mm, respectively.

An image processing strategy and stimulation method are used to convert the target spots and gratings into a pattern of electrode stimulation (Figure 2a). The pattern is represented as neural activation values at a set of discrete locations across the retina that we will refer to as pixels. A simulation or animal experiment is then performed and the resulting spatial pattern of neural activation is measured. The pattern is repeated over spots at 16 different randomized locations within a central 1000 μm × 1000 μm location and for gratings using 16 different random phases and orientations. These patterns are registered to each other so that the mean neural activation value across all 16 patterns can be calculated and a single final evoked neural activation pattern calculated. This repetition and averaging performs two functions:

1. It allows for statistical smoothing that avoids any given measurement from being overly influenced by the particular relative position of the electrodes and the target pattern.
2. It can be interpreted as a highly simplified model of the normal cognitive integration undertaken using saccades or head movements.

This average evoked pattern is considered to be the final pattern that can be compared to the target. This comparison is done using mean squared error difference for target spots, *ε*_TS_, and target gratings, *ε*_TG_, on a pixel-by-pixel basis (Figure 2b).

The evoked pattern is *also* compared to a set of reference spots with adjacent positions to the target spots, *ε*_RS,*n*_, or reference gratings with different orientations, *ε*_RG,*n*_ (Figure 2c). The reference gratings are chosen to be oriented at 30°, -30°, 60°, -60°, and 90° relative to the target grating (Figure 2c, left). The reference spots are placed at a distance of 0.5 cycles from the target spot at equally spaced 30°, -30°, 90°, -90°, 150°, and -150° positions (Figure 2c, right), where 1 cycle refers to the spatial period of a grating stimulus at the equivalent resolution level.

The resulting mean squared errors are used to calculate the mean and standard deviation of *ε*_R,*n*_, which are then used to calculate a perceptible difference, *d*′, for each comparison,

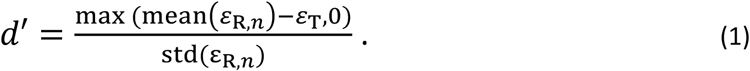

This has been modified to recognise that we know that mean (*ε*_R,*n*_) > *ε*_T_ because *ε*_R,*n*_ are errors based on reference patterns that were not used to create the evoked pattern of neural activity. The *d*′ measurement is repeated with varying spot sizes and grating periods. These feature sizes are varied to be six logarithmically spaced values between a minimum of 0.25 times the electrode pitch and a maximum of 1000 μm. This maximum value was chosen based on the size of our simulated electrode array (5500 μm × 3400 μm, see Vision Model below).

Finally, the perceptible feature size is calculated as the feature size with a *d*′ value equal to 3 using linear interpolation between each of the six feature sizes. This value was chosen because it represents 3 standard deviations, which is associated with an error rate of 0.3 %.

The parameters used in the acuity measurement are shown in Table 1.

**Table 1:** Summary of the parameters associated with the acuity measurement.

| Parameter | Value |
| --- | --- |
| Number of Feature Sizes Used | 6 |
| Minimum / Maximum Feature Size | 0.25×Pitch / 1000 μm |
| Number of Jittered Spots / Gratings | 16 / 16 |
| Range of Random Target Location | 1000 μm × 1000 μm |
| Reference Spot Distance from Target Spot | 0.5 × Feature Size |
| Five Reference Grating Angles | 30°, -30°, 60°, -60°, 90° |
| Six Reference Spot Angles | 30°, -30°, 90°, -90°, 150°, -150° |
| Perceptible $d'$ | 3 |

### 2.2 Acuity Measurement Confirmation

Conventional clinical visual acuity tests use optotypes to assess visual acuity. These are also used in this investigation to provide a way to assess the calculated visual acuity level. A Landolt-C is used, with a gap-width 1/5 of the overall letter width, and a letter E with features of 1/5 the overall letter width. In each case, letters are displayed at the measured size that should allow perception of the orientation of the letter, as well as -20% and +20% this size.

### 2.3 Neural Simulations and Stimulation Strategies

Although the acuity measurement is intended to apply to any simulation or experimental measurement of neural activity, we use simulations of neural activation that require a vision model in this investigation. Given that the acuity measure is explicitly proposed for scenarios with overlapping regions of neural activation, we compare a stimulation strategy that accounts for overlapping regions of neural activation with one that does not.

#### 2.3.1 Vision Model

A computational model converts patterns of electrode settings into patterns of neural activation. The model used here has been adapted from a model that is experimentally verified to be accurate [12] and is applicable to retinal implants as a model of either retinal or cortical activity and to cortical implants as a model of cortical activity [11]. The model is not intended to be assessed in this investigation or be realistic in every detail; it is intended to be used to demonstrate the features of the acuity measure.

The model is identical to the model used in previous publications and is described by [15,19]. For clarity, we provide a short overview of the model here. The model uses scaled units with S – spikes, T – time, I – electric current, L – length. Briefly, a linear non-linear model converts electrode settings 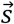 [I] into a pattern of neural activation 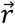 [ST^-1^],

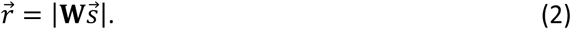

The columns of the matrix **W** contain the spread of neural activity that results from activation of each electrode. Here, for simplicity, these spreads are modelled as circularly symmetric Gaussian functions with a spread value (standard deviation) of σ. This uniform approach allows straightforward comparison between implants with spreads of different sizes. The electrode array was modelled as a total width of 5500 μm and height of 3400 μm. This corresponds to a visual field of approximately 19°×12°.

#### 2.3.2 Conventional Strategy

The conventional stimulation strategy assumes that each electrode should be activated in proportion to the amplitude of the desired neural activity pattern at its location. In this study, this is implemented by projecting spread of neural activation of each electrode onto the target pattern,

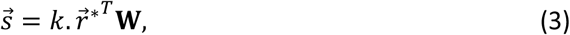

where 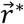 is a target neural activation pattern and *k* is a normalization constant chosen to correct the overall activation level of the array [15]. Safe electrode stimulation amplitudes are enforced by multiplicatively scaling all stimulation amplitudes if any electrode is above the safe limit.

#### 2.3.3 Neural Activity Shaping (NAS) strategy

An alternative approach, developed by our group, uses an inverse model to manage the overlapping spreads of neural activation [15,19]. The target neural activation pattern is converted into appropriate electrode settings as described in [15]. This inverse model is based on targeting only positive values of the 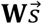 term in Equation (2) so we can assume that 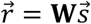. It is assumed that we wish to minimize the mean squared error between the evoked and target patterns of neural activity,

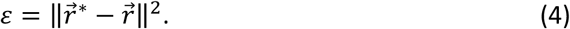

Substituting the solution 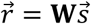 gives the objective function,

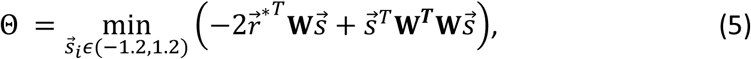

using maximum and minimum electrode settings of magnitude 1.2. This can be solved for the elements of 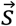 using a quadratic programming approach, as described in [15], and includes an explicit limit to enforce safe electrode stimulation levels. In the present investigation, we use all eigenvalues in the singular value decomposition of **W**.

#### 2.3.4 Safe Electrode Settings

A feature of the NAS strategy is that the results are influenced by the maximum safe electrode amplitudes. For the simulations used in this study, the maximum safe electrode amplitude was set to 1.2 [I]. This is 20% higher than the level required for a single electrode activation to induce maximum neural activation of 1 [I].

## 3 Results

A simulation is created of an evoke neural activity patten with an electrode array with an electrode pitch of 450 μm and neural activity spreads of *σ* = 450 μm (Figure 3a). This is an array that results in overlaps in the spread of neural activity, which means that its acuity cannot be estimated by simply assuming that the acuity is determined by the electrode pitch, which gives a value of 94 MAR.

**Figure 3:**
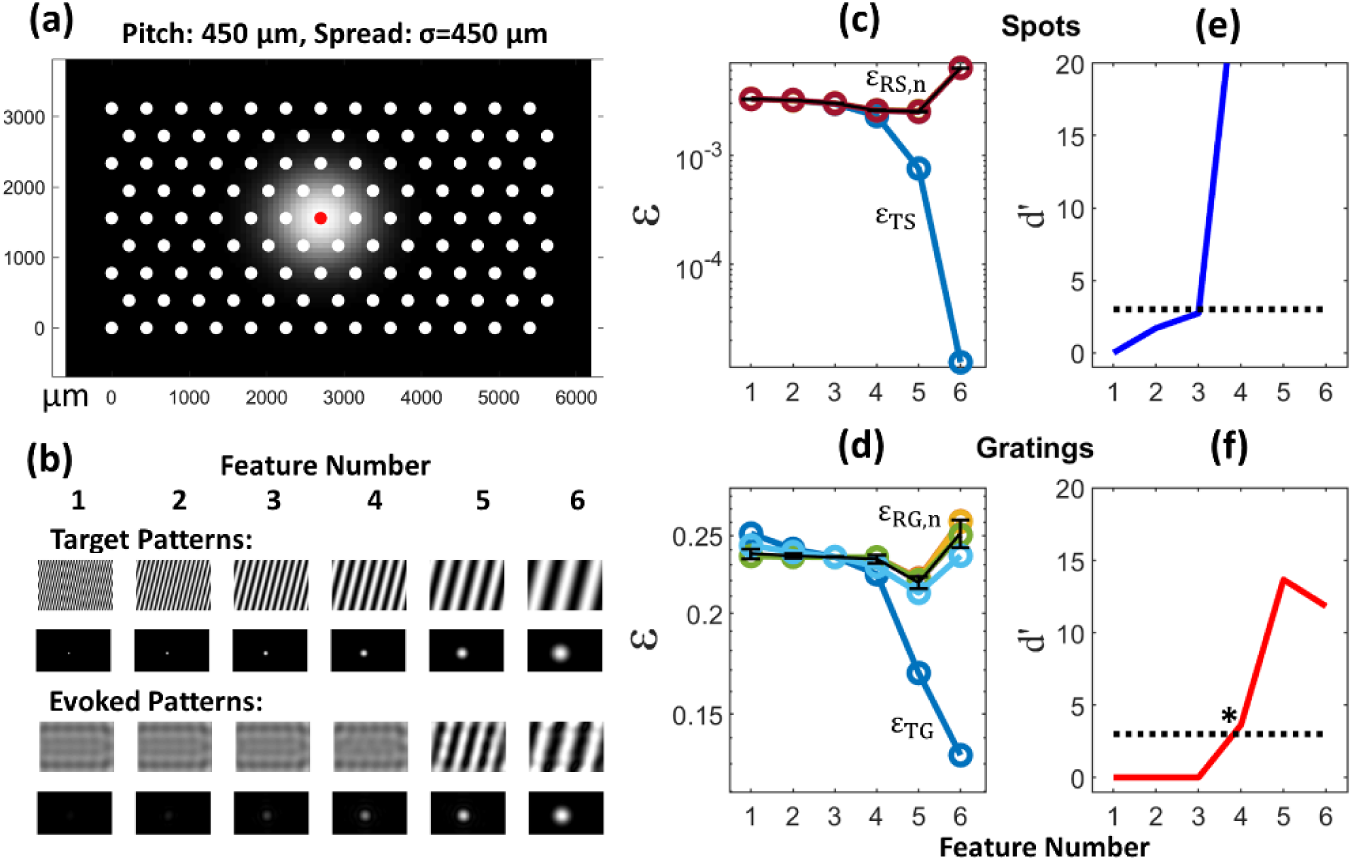
Assessment of the d′ value of an implant with overlapping spreads of neural activity using the NAS strategy. Electrode pitch of 450 μm and neural activity spreads of σ = 450 μm. (a) The implant configuration to be assessed. (b) An example set of targets used to calculate d′ values and the resulting evoked patterns. (c) and (d) The particular MSE values for the targets (ε_TS_ and ε_TG_) and reference patterns (ε_RS,n_ and ε_RG,n_). The means and standard deviations associated with the reference patterns are shown in black. (e) and (f) The d′ values calculated from these MSE values using Equation 1. The dotted line indicates the d^′^ = 3 cut-off value. The asterisk in (f) indicates the smallest feature size at which the reference and target patterns are distinguishable for both spots and gratings and constitutes the estimated clinical visual acuity.

A particular set of targets and evoked activities is shown for the array (Figure 3b) using the NAS strategy with the electrode current safety limit. The *d*′ values for these particular features are calculated as described in Methods using the mean squared errors between these evoked activities and a set of reference activities (Figure 3c and 3d). A range of feature sizes are used with a range of dot positions and grating orientations and phases, and a *d*′ value is calculated for each (Figure 3e and 3f).

The grating *d*′ curve intersects a value of *d*′ = 3 between the 3^rd^ and 4^th^ feature size (Figure 3f) which is larger than the value estimated using spots (Figure 3e). Using linear interpolation, the precise crossing value corresponds to a visual acuity of 55 MAR.

Different arrays with the same pitch of 450 μm used in the foregoing analysis are simulated and each array’s representation of a hand, held at arm’s length, is shown along with the outcome of this measurement in the form of appropriately scaled optotypes (Figure 4). This post-hoc qualitative assessment is shown to provide assurance that the results of this proposed acuity measure are reasonable. This process if completed for neural activity spread of *σ* = 100 μm using the conventional stimulation or NAS stimulation strategy, which with isolated phosphenes give the same results (Figure 4a); with a neural activity spread of *σ* = 450 μm using the conventional strategy (Figure 4b); and with a neural activity spread of *σ* = 450 μm using the NAS strategy (Figure 4c).

**Figure 4:**
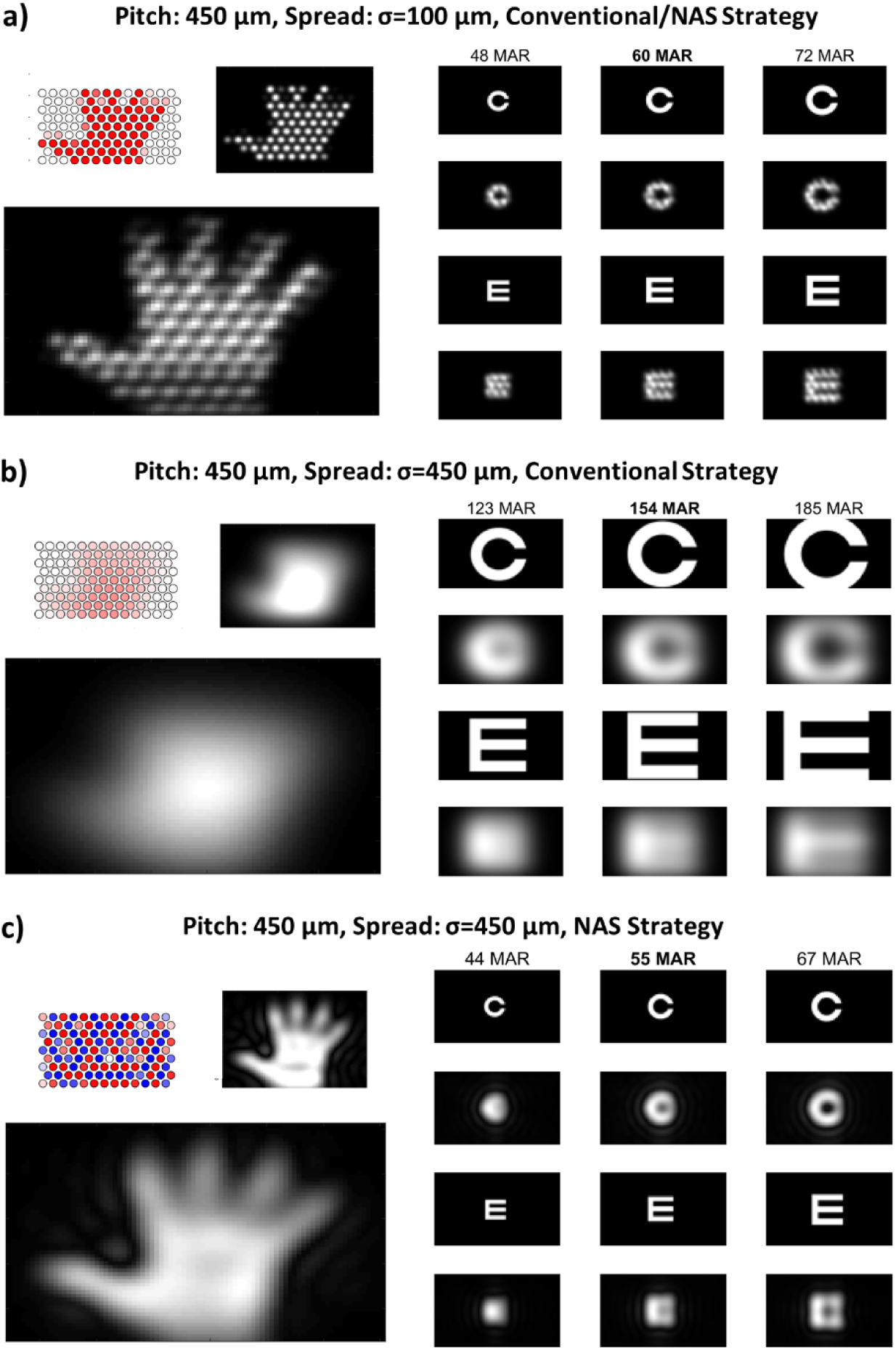
Example qualitative acuity measurement results for implants all with an electrode pitch of 450 μm, corresponding to an acuity of 60 MAR under the conventional assumptions based only on electrode pitch used in [1,4,9]. The left column shows the results for a hand at arm’s length for each implant with the electrode and neural pattern for an individual stimulus shown above each larger image. These show the neural patterns after integration over 16 registered trials. The right column shows optotypes scaled to be -20% (left) the same as (middle) and +20% (right) of the acuity level assessed using the proposed method. The first and third rows show target patterns; the second and fourth rows show corresponding resultant neural activity patterns. (a) Activity spread σ = 100 μm using the conventional strategy. (b) Activity spread σ = 450 μm using the conventional strategy. (c) Activity spread σ = 450 μm using the NAS strategy.

By examining the three optotype scales, it is possible to see that at the lower level (−20% of the estimated acuity value) it is very difficult to perceive the orientation of the letter using the evoked activity pattern. At the upper level (+20% of the estimated acuity value), it is straightforward to identify the orientations of the optotypes. This confirms that the acuity measurement is providing a useful assessment of the visual acuity across both isolated and overlapping spreads of neural activity.

The method for assessing visual acuity is used to assess the acuity of electrode arrays with a range of electrode pitches and isolated and overlapping spreads of neural activity (Figure 5). In cases where the spread of neural activation from each electrode is isolated and non-overlapping, the conventional strategy and NAS strategy give identical results (Figure 5a). The conventional method of estimating acuity assumes that acuity is proportional to the electrode pitch [1,4,9]. In this case, visual features of a size proportional to half the pitch can be perceived (the black line in Figure 5a). However, in cases where there is overlapping neural activation, the straightforward method gives highly inaccurate estimates of visual acuity when using a conventional stimulation strategy. Compare the blue curve and the black line in Figure 5b; it is apparent that a better estimate would be proportional to the size of the phosphene spread (the dashed black line).

**Figure 5:**
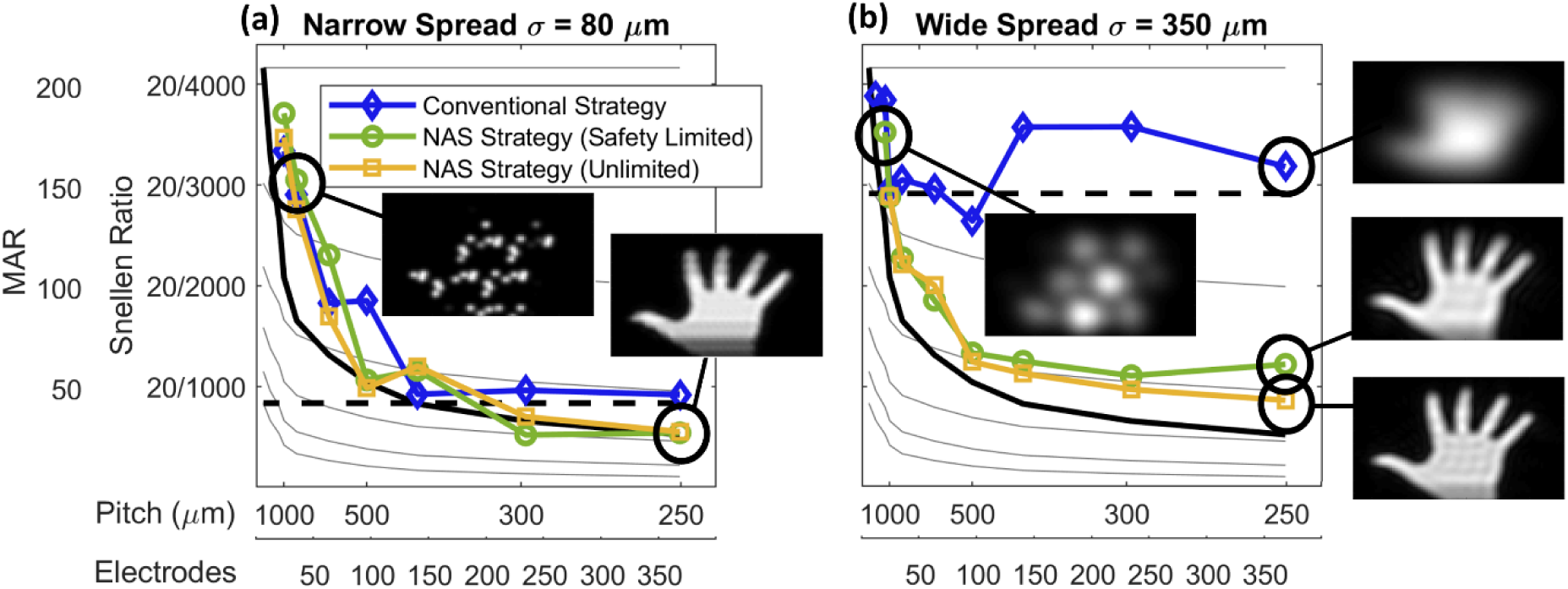
Acuity with varying electrode pitch and stimulation strategy under (a) narrow spread (isolated non-overlapping) and (b) wide spread (overlapping) neural activity conditions. The blue lines (diamond symbols) show the visual acuity at each electrode configuration using the conventional stimulation strategy, green line (circle symbol) uses the NAS strategy with an electrode safety limit, and the yellow line (square symbol) uses the NAS strategy without any electrode safety limit. The black solid lines show theoretical maximum acuity due to the electrode pitch, corresponding to half the electrode pitch. The dashed black lines show the apparent limit due to neural activity spread (2×σ). The black circles indicate data points for which an image of evoked activity is provided for a hand held at arm’s length.

In cases with overlapping spreads of neural activity, the NAS strategy can initially track the maximum possible acuity provided by the electrode pitch; compare the blue curve and the green/yellow curves with the black line in Figure 5b. When a safety limit is not placed on the electrodes, the simulation shows that, in theory, the NAS strategy could continue to track this level of acuity because the algorithm can manipulate the pattern of activation to an arbitrarily high degree. However, in practice, it is necessary to use safe electrode limits. Under the NAS strategy, this limits the capacity of the algorithm to manipulate the pattern of activation and the acuity plateaus (green curves in Figure 5).

## 4 Discussion

This investigation demonstrates a method for assessing the acuity provided by visual prostheses in a preclinical setting. The method uses patterns of evoked neural activation based on target and reference patterns of spots and gratings. These evoked patterns can be obtained from recordings obtained from the retina or cortex in animal experiments or from computational simulations of neural activation. By examining qualitative images of standard clinical optotypes (letters and hands) scaled based on the results of the proposed acuity measurement, it is seen that the results are reliable in providing a reasonable estimate of true visual acuity.

The method is intended to use as few recordings as possible so that data recording requirements are minimised. However, parameters of the method can be adjusted to decrease the data requirements (and measurement time) further or, alternatively, to increase the accuracy of the results. This could be achieved by changing the number of feature sizes tested (six were used in this investigation) and/or changing the number of repetitions of the target (a total of 16 repetitions were used in this investigation).

Implicit in the proposed method is the idea that contrast in the levels of activation within evoked patterns of neural activity are a reasonable estimate of the contrast in the levels of brightness within patterns of visual perception. While it is known that there is not a one-to-one match between these quantities, there is sufficient evidence to make this a useful pre-clinical measure of visual perception [20].

It was important to make the approach as similar as possible to the clinical approach so the method proposes the use of both spots and gratings as visual targets because both are used in clinical settings in assessment of visual acuity levels. In practice, it was observed that gratings were more easily perceived than spots when the evoked neural activity was isolated and non-overlapping, while spots were more easily perceived than gratings when there were wide, overlapping spreads of neural activity. Combining both in a single measure of acuity takes advantage of both measures of visual perception.

The method proposed in this investigation includes a simple model of cognitive integration by repeating each target presentation 16 times before calculating the mean squared error values. This is a simple way to capture the fact that, in a clinical setting, someone may scan their vision across the pattern to utilise different parts of their array to represent the information. Alternative, more explicit models of cognitive integration may also be developed to capture the effects of visual saccades and scanning [21].

